# Cardiac metabolic effects of K_Na_1.2 channel deletion, and evidence for its mitochondrial localization

**DOI:** 10.1101/223321

**Authors:** Charles O. Smith, Yves T. Wang, Sergiy M Nadtochiy, James H. Miller, Elizabeth A. Jonas, Robert T. Dirksen, Keith Nehrke, Paul S. Brookes

## Abstract

Controversy surrounds the molecular identity of mitochondrial K^+^ channels that are important for protection against cardiac ischemia-reperfusion injury. While K_Na_1.2 (*Kcnt2* gene) is necessary for cardioprotection by volatile anesthetics, electrophysiologic evidence for a channel of this type in mitochondria is lacking. The endogenous physiologic role of a potential mito-K_Na_1.2 channel is also unclear. Herein, single channel patch-clamp of 27 independent cardiac mitochondrial inner membrane (mitoplast) preparations from wild type (WT) mice yielded 6 channels matching the known ion-sensitivity, ion-selectivity, pharmacology and conductance properties of K_Na_1.2 (slope conductance 138±1 pS). However, similar experiments on 40 preparations from *Kcnt2*^-/-^ mice yielded zero such channels. The K_Na_ opener bithionol uncoupled respiration in WT but not *Kcnt2*^-/-^ cardiomyocytes. Furthermore, when oxidizing only fat as substrate, *Kcnt2*^-/-^ cardiomyocytes and hearts were less responsive to increases in energetic demand. *Kcnt2*^-/-^ mice also had elevated body fat, but no baseline differences in the cardiac metabolome. These data support the existence of a cardiac mitochondrial K_Na_1.2 channel, and a role for cardiac K_Na_1.2 in regulating metabolism under conditions of high energetic demand.

## Introduction

Numerous strategies for protection of the heart and other organs against ischemia-reperfusion (IR) injury are thought to require activation of K^+^ channels in the mitochondrial inner membrane (for review see (1)). This includes ischemic preconditioning (IPC), volatile anesthetic preconditioning (APC), and pharmacologic cardioprotection by K^+^ channel activators such as NS-11021, bithionol and diazoxide(2-4). Concurrently, several K^+^ channels have been reported in mitochondria including: ATP activated (K_ATP_)(5), small conductance Ca^2+^ activated(SK)(6, 7),and splice variants of large conductance Ca^2+^ activated(BK)(8, 9). However, in only a small number of cases has the molecular(genetic)identity of specific mitochondrial channels involved in cardioprotection been proposed(3, 9-12). No K^+^ channels exist in the “MitoCarta” database of verified mitochondrial proteins(13)and no canonical mitochondrial target sequences have been identified in any K^+^ channel genes(1). As such, mystery surrounds the mechanism(s) of K^+^ channel targeting to mitochondria.

Mammalian Na^+^-activated-K^+^ (K_Na_) channels are encoded by two genes: *Kcnt1* (14) and *Kcnt2* (15), which produce the K_Na_1.1 (Slack/SLO2.2) and K_Na_1.2 (Slick/SLO2.1) channels respectively. Both K_Na_1.1 and K_Na_1.2 channels play important neurologic roles in the termination of seizure progression in epilepsy(16, 17). Although K_Na_1.2 expression has been reported in cardiac tissue(15, 18),and a generic K_Na_ channel activity has been demonstrated in the cardiac cell membrane(19),notably the *Kcnt2^-/-^* mice have no cardiac phenotype(3, 18). Thus relatively little is known regarding the physiologic role of K_Na_1.2 channels in the heart, including their subcellular location. Previously we showed that K_Na_1.2 is essential for cardiac APC, with hearts from *Kcnt2*^-/-^ mice incapable of being protected against IR injury by isoflurane(3). Additionally, we showed that the K_Na_ opener bithionol (BT) is cardioprotective in WT mice but not *Kcnt2*^-/-^ mice (3, 20). Further, both BT and isoflurane activated a K^+^ flux in cardiac mitochondria isolated from WT mice, but not those from *Kcnt2*^-/-^ mice(3). These observations led us to hypothesize that a K_Na_1.2 channel may exist in cardiac mitochondria.

There is considerable evidence that mitochondrial K^+^ channel activity can lessen the impact of IR injury(1). less is known about the endogenous physiologic role(s) of mitochondrial K^+^ channels(21),and in particular mitochondrial K_Na_ channels. Na^+^ enters mitochondria predominantly via a Na^+^/Ca^2+^ exchanger (NCLX) or a Na^+^/monocarboxylate transporter (MCT), with minor contribution from a Na^+^/H^+^ exchanger(NHE)(22). Under conditions of elevated mitochondrial Na^+^ uptake (Ca^2+^ overload or cytosolic Na^+^ overload driving excessive NCLX activity, or increased cytosolic Na^+^-lactate driving Na^+^ uptake via the MCT), matrix Na^+^ may attain levels that can activate K_Na_ channels (23-25). K^+^ influx via the K_Na_ would dissipate mitochondrial membrane potential, thereby relieving a driving force both for Ca^2+^ uptake and NCLX activity. In effect, a mitochondrial K_Na_ channel acts as a “safety valve” to prevent ionic imbalance bought about by excessive Na^+^ uptake.

Herein, using electrophysiologic techniques (mitoplast patch-clamp) we demonstrate that mitochondria contain a K^+^ channel that matches the known ion-sensitivity, ion-selectivity, pharmacology and conductance properties of K_Na_1.2. Bioenergetic studies of WT and *Kcnt2^-/-^* hearts and cardiomyocytes also reveal a potential role for cardiac K_Na_1.2 in the metabolic response to high energetic demand.

## Materials and Methods

### Animals

Male and female mice were housed in an AAALAC-accredited pathogen-free facility with water and food available *ad libitum*. All procedures were locally approved and in accordance with the NIH *Guide for the Care and Use of Laboratory Animals* (2011 revision). Mice were on a C57BL/6J background for >6 generations and periodically backcrossed to fresh stocks. Mice were bred from *Kcnt2*^+/-^ parents, and male and females were separated but littermate WT and *Kcnt2*^-/-^ progeny were maintained in the same cages. Mice were genotyped by tail-clip PCR (Figure 1A), with DNA extraction by a Qiagen DNeasy^™^Kit (Hilden, Germany) and genotyping by a Kapa Biosystems KAPA2G Kit (Wilmington, MA). Primers used were (5’ 3’) forward-AGGCAGCCATAGCTTTAGAGA and reverse CTCCTCATCGTGTGGTCCTA, yielding amplicons at 822 and 547 bp for WT and *Kcnt2^-/-^* respectively. Due to the same personnel handling mice and performing experiments, studies were not blinded to genotype.

**Figure 1:**
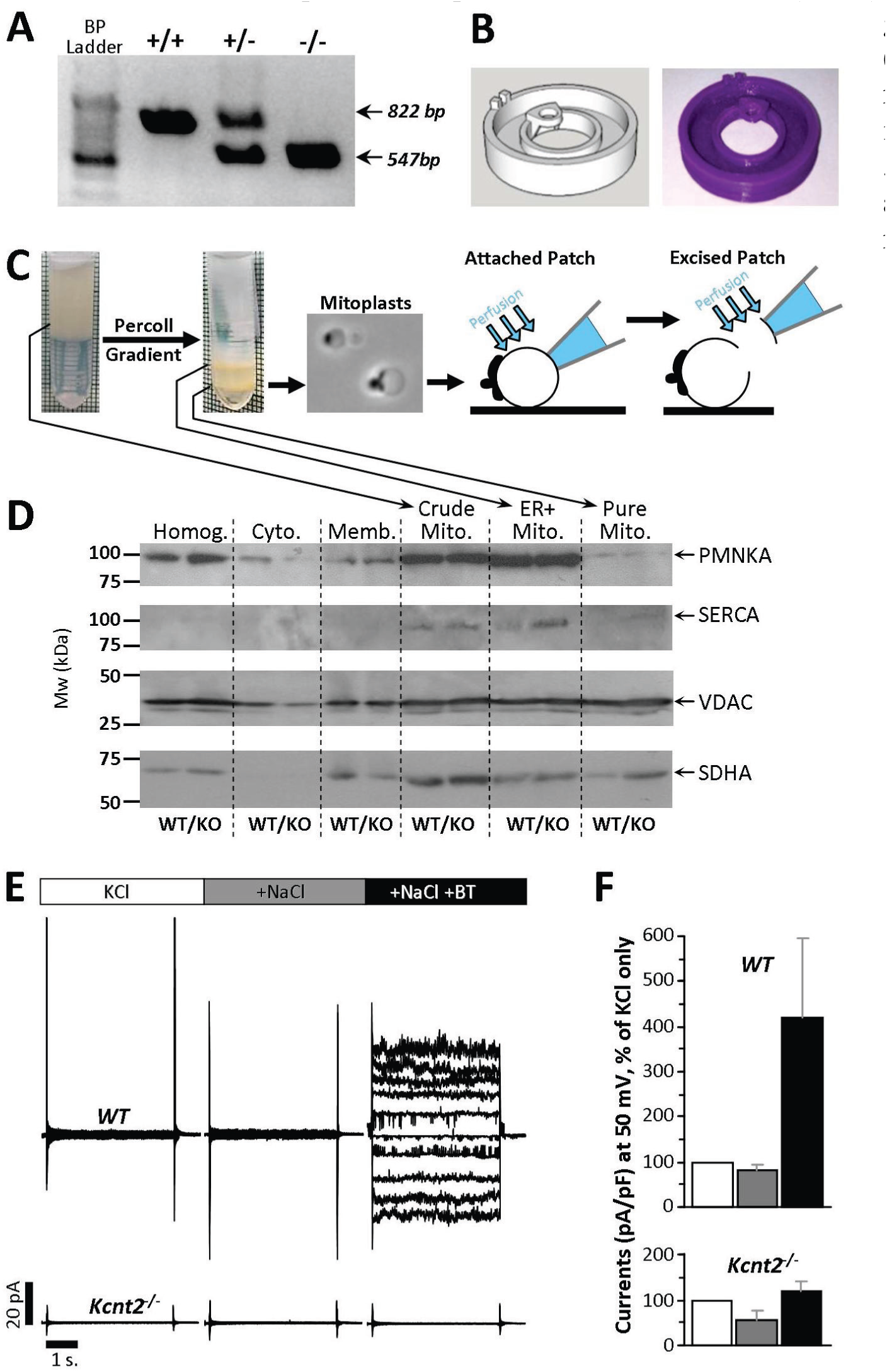
mice, mitoplast purification and attached patch clamp. **(A)** PCR analysis of tail clip genotyping of WT, heterozygous, and *Kcnt2^-/-^*, mice. 822 bp amplicon expected for WT allele and 547 bp expected for knockout allele. **(B)** Custom 3D printed micro chamber for patch-clamp of mitoplasts with computer model (left) and final product (right). Stereolithography file is deposited at: https://3dprint.nih.gov/discover/3dpx-008253. **(C)** Schematic depicting mitochondrial purification, mitoplast preparation, attached patch and excised patch configuration. **(D)** Western blot of proteins from different cellular fractions during mitochondrial purification. (Homog: homogenate, Cyto: cytosol, Memb: crude membrane, Crude Mito: mitochondrial enriched fraction, ER+Mito: upper band following Percoll^™^, Pure Mito: lower band following Percoll^™^). PMNKA: plasma membrane Na+/K+-ATPase, SERCA: sarco-endoplasmic reticulum Ca2+-ATPase, VDAC: mitochondrial outer membrane voltage dependent anion channel, SDHA: mitochondrial inner membrane succinate dehydrogenase subunit A. **(E)** Representative recordings from attached patch clamp of WT and *Kcnt2*^-/-^ mitoplasts, with perfusion of 150 mM KCl (white), after addition of 40 mM NaCl (dark grey), and after addition of 10 µM BT (Black). **(F)** Quantitation of traces normalized to pA/pF and shown as % increase in current at holding potential of 50mV (Same color scheme as **(E)**). Data are means ± SD, from 4-10 independent mitoplast preparations.

### Patch Clamp of Mitochondrial Inner Membranes (Mitoplasts)

Following anesthesia (tribromoethanol 200mg/kg ip) the heart from one 8-12 week old mouse was rapidly excised, washed and chopped in ice-cold mitochondrial isolation medium (MIM, in mM: 300 sucrose, 20 Tris, 2 EGTA, pH 7.35 at 4 °C). All steps were performed on ice. Tissue was homogenized (Tissumizer^TM^, IKA Inc., Wilmington NC) then centrifuged at 700 × *g*, 5 min. Supernatants were saved and pellets re-homogenized and re-centrifuged. Pooled supernatants were then centrifuged at 10,000 × *g*, 10 min. The crude mitochondrial pellet was suspended in 0.2 ml MIM and layered over 1.75 ml of 30% osmotically-balanced Percoll^™^, in a round-bottomed microcentrifuge tube, and centrifuged at 14,000 × *g*, 1 hr. Two mitochondrial layers were apparent (Figure 1C), of which the lower (purified mitochondria) was washed twice by centrifugation. The mitochondrial pellet (~25 µl) was suspended in 0.5 ml swelling buffer (in mM: 30 KCl, 20 HEPES, 1 EGTA, pH 7.2) for 15 min. Centrifugation (1,000 × *g*, 30 s.) afforded a mitoplast pellet, resuspended in ~20 µl MIM for immediate use in patch-clamp studies (N=27 WT, 40 *Kcnt2*^-/-^).

For mitoplast attached patch-clamp studies, mitoplasts were diluted 1:100 in a bath solution (in mM: 150 KCl, 20 HEPES, 1 EGTA, pH 7.2) and a 10 µl drop was placed on a glass coverslip attached to a custom 3D printed micro-chamber (Figure 1B). Electrodes (40-100 M) (Sutter Instruments, Novato CA) were filled with pipette solution (in mM: 150 KCl, 0.025 NaCl, 20 HEPES, 1 EGTA, pH 7.2). The bath was exchanged with buffer additionally containing 40 mM NaCl, or 40mM NaCl and 10 µM bithionol (BT, from stock in DMSO, final DMSO < 0.01% v/v). For excised patch experiments mitoplasts were diluted 1:100 in patch seal buffer (in mM: 60 KCl, 80 K-gluconate, 40 LiCl, 0.025 NaCl, 0.1 CaCl_2_(calculated free), 20 HEPES, 1 EGTA, pH 7.2) Electrodes were filled with pipette solution (in mM: 125 KCl, 15 K-gluconate, 15 LiCl 0.025 NaCl, 20 HEPES, 1 EGTA). Mitoplasts were identified by their round shape and presence of a “cap” structure (Figure 1C). After formation of G seals, patches were excised and inside-out currents were recorded using an Axopatch 200B amplifier and Clampex10 software (Molecular Devices, Sunnyvale CA). All holding potentials reported are those applied to the patch pipette interior. The electrical connection was made using Ag/AgCl electrodes and an agar 2M KCl salt bridge at the ground electrode. (Note: not all channels yielded currents at all potentials, and seal integrity was often compromised at the extremes of this range). Data was digitized and recorded at 10 kHz and filtered using an 8-pole low pass 2 kHz filter. Patches were recorded under flow (0.1 ml/min.) of: (i) Ca^2+^ free patch seal buffer with 0.076 mM sucrose for osmotic balance, (ii) as above, with LiCl replaced with 40 mM NaCl, (iii) further addition of 2.5 µM BT (from stock in DMSO, final DMSO < 0.01% v/v). Although the study of K^+^ channels typically employs gluconate salts to exclude the possibility that measured conductances are due to Cl^-^ channels, Cl^-^ salts were used herein because K_Na_1.2 activity is strongly enhanced in the presence of Cl^-^(15). All buffers were filtered (0.22 µm) immediately before use. Single channel analysis was performed using Clampfit 10.0 single channel search (Molecular Devices). Dwell times were calculated using “single-channel search” using threshold crossing.

### Cardiomyocyte Isolation and Respiration Measurements

Mouse primary adult ventricular cardiomyocytes were isolated by collagenase perfusion as previously described(3). Cells were step-wise rendered tolerant to 1.8 mM Ca^2+^, and the final pellet suspended in 1 ml MEM (GIBCO cat # 11095-080, supplemented with 1.8 mM, CaCl_2_ 2.5% FBS and pen/strep). Cell viability and yield were determined using Trypan blue and a hemocytometer. Only preparations with >85% viable rod-shaped cells were used for experiments. Cells were seeded at 2000/well on Seahorse^™^ XF96 V3-PS plates (Agilent, Billerica MA) and equilibrated for 1 hr. MEM was replaced with unbuffered DMEM (pH 7.4) containing various carbon sources (in mM: 5 glucose, 0.1 palmitate, 4 glutamine, 5 galactose, 5 lactate, 1 pyruvate) and either 10 mM 2-deoxyglucose or 20 µM etomixir as detailed in results. All conditions with palmitate had 0.1 mM L-carnitine. Oxygen consumption rates (OCR) were measured using an XF96 extracellular flux analyzer.

### Ex-vivo Heart Perfusion

Mouse hearts were perfused in constant flow(4ml/min)Langendorff mode as previously described(3). Krebs-Henseleit buffer (KH, in mM: 118 NaCl, 4.7 KCl, 25 NaHCO_3_, 1.2 MgSO_4_, 1.2 KH_2_PO_4_, and 2.5 CaCl_2_, gassed with 95/5 O_2_/CO_2_, 37 °C) was supplemented with either 5 mM glucose, or 0.1 mM BSA-conjugated palmitate. Left ventricular pressure was measured via a water-filled transducer-linked left ventricular balloon. Left ventricular and coronary root pressures were monitored and digitally recorded at 1 kHz (DATAQ, Akron OH). After equilibration hearts were treated with isoproterenol (100 nM final) for 5 min.

### Electron Microscopy

Hearts were fixed in 4 % paraformldehyde + 2.5 % glutaraldehyde in Millonig’s phosphate buffer (0.2 M NaH_2_PO_4_/Na_2_HPO_4_, 0.5 % NaCl, pH 7.4). 1 mm cubes were processed and digitally photographed on a Hitachi 7650 electron microscope. Analysis of images was performed using NIH ImageJ software. Mitochondrial areas and density were placed in to 11 or 13 bins respectively and the resulting histograms were fitted to a single Gaussian. Form-factor was calculated as 1/((4.area)/(perimeter^2^)) and aspect-ratio was calculated as (major axis/minor axis).

### Metabolomics

WT and *Kcnt2*^-/-^ hearts (N=7 per group) were perfused as above in KH buffer supplemented with glucose plus palmitate for 20 min., then freeze-clamped with Wollenberger tongs in liquid N_2_ and ground to powder. Samples representing ~50% of each heart (50 mg) were shipped to Metabolon Inc. (Research Triangle Park, NC) on dry ice, extracted by standard procedures, and analyzed by LC-MS/MS and GC-MS/MS (Metabolon “Global Metabolomics” solution) to measure the relative steady-state abundance of metabolites.

Data for each run were median-normalized. Overall, 527 metabolites were identified, of which 26 (4.9%) were removed due to insufficient replicates, yielding 7014 theoretical individual data points (501 × N=7 × 2 groups). A further 229 outliers (>1 standard deviation from the mean) were removed, representing 3.3 % of the data. Missing values were imputed as weighted medians(26). Metabolomic data were analyzed using free Metaboanalyst software(27). In a separate series of experiments, WT and *Kcnt2*^-/-^ hearts were perfused in KH buffer supplemented with fat as the only carbon source, and adenine nucleotide levels (ATP, ADP, AMP) were measured as previously described(28). Energy charge was calculated as (ATP+½ADP/(ATP+ADP+AMP).

### Immunoblotting

Sample protein was determined by the Folin-Phenol (Lowry) assay. Non-mitochondrial samples were diluted 2x in Laemmli sample loading buffer (SLB) and incubated at 95 °C for 1 min., while mitochondrial samples were diluted 2x in SLB containing 5x the standard concentration of SDS and incubated at 25 °C for 30 min. Samples were separated by SDS-PAGE (10% gels) and transferred to nitrocellulose, followed by probing with antibodies as recommended by manufacturer protocols. Detection employed HRP-linked secondary antibodies with enhanced chemiluimnescence (GE Biosciences). Developed ECL film images were quantified by densitometry using NIH ImageJ software (N=3-4 mice per genotype).

### Body Composition, Fasting Glucose Response, and Electrocardiogram

84 day old(12 week)WT and littermate *Kcnt2*^-/-^ male mice were anesthetized as described above, and body fat content measured using dual energy X-ray absorptometry (DEXA) scanning (Lunar PIXImus densitometer, GE, Fitchburg WI). Blood glucose was measured using a True2Go^™^ glucose meter with TrueTest^™^ glucose strips (Trividia Health, Fort Lauderdale FL). Mice were fasted overnight in cleaned cages with access to water and cotton bedding. Alternatively, after anesthesia mice electrocardiograms were recorded using a three electrode EKG amplifier (Harvard Apparatus, Cambridge MA). EKGs were averaged for each animal from ten different segments of the trace, each containing R_1_-S_1_-T_1_-P_2_-Q_2_-R_2_ waves.

### qPCR analysis

mRNA was extracted from heart homogenates with acid phenol/TRIzol according to the Direct-zol RNA MiniPrep Kit R2050 (Zymo Research, Irvine CA) as described(29). cDNAs were prepared using an iScript kit (170-8891, BioRad). qPCR analysis was performed using a BioRad PrimePCR^TM^ “Regulation of lipid metabolism-PPAR” M96 Predesigned 96-well panel for use with SYBR^®^ Green (Cat # 10031585).

### Replicates & Statistics

Numbers of individual replicates for experiments are listed in each figure legend. For samples comparing WT and *Kcnt2*^-/-^, one “N” equals one animal (i.e., biological replicates). Statistical differences between WT and *Kcnt2*^-/-^ were determined using two-way ANOVA with a Bonferroni correction for multiple testing, followed by post-hoc paired or non-paired *t*-tests (p<0.05 cut-off).

## Results

### Kcnt2 knockout

Figure 1A shows the result of a genotyping experiment on WT and *Kcnt2*^-/-^ mice, indicating the expected amplicons (see methods). The deletion targets exon 22 of the protein, and results in no residual protein in the knockouts(18), as we have also previously confirmed in the heart(3).

### Cardiac Mitochondria Contain a K_Na_1.2 Channel

To investigate cardiac mitochondrial K^+^ channels, we performed electrophysiology studies on Percoll^™^-purified isolated mitochondrial inner membranes (mitoplasts) from hearts of wild type(WT)or *Kcnt2*^-/-^ mice (schematic, Figure 1C). Mitochondrial enrichment was verified by western blotting for mitochondrial proteins, and purity was confirmed by western blotting for non-mitochondrial membrane proteins (Figure 1D). Mitoplasts were readily identified under light microscopy (Figure 1C) and high resistance seals (2-10 G) were formed with the inner membrane opposite the outer membrane cap. In a preliminary series of experiments using attached mitoplast patch configuration, channel activity in WT was not activated by NaCl alone, but manifested upon further addition of the K_Na_ activator bithionol(BT) (3). No activation by NaCl or BT was seen in *Kcnt2^-/-^* mitoplasts (Figure 1E/F). The absence of activation by NaCl alone in WT mitoplasts suggested that the channel’s ion-sensing site may be on the matrix side of the inner membrane, and thus inaccessible to Na^+^ from the bath perfusion on this time scale. To investigate such a channel, excised mitoplast patch was employed, affording access to the matrix side of the inner membrane (Figure 1C).

Channel activity was observed in a high proportion of independent preparations obtained from both WT (85%) and *Kcnt2*^-/-^ (76%) hearts (Figure 2A), consistent with previous reports that mitochondria contain numerous K^+^ channels (1, 30). A multiple level screen was performed to triage recordings containing channels other than K_Na_1.2 (Figure 2B-D). The bath recording solution was sequentially switched from initial (containing 100 µM CaCl_2_), to 40 mM LiCl, then 40 mM NaCl, and finally 40 mM NaCl plus 2.5 µM BT. Under each condition, channel activity was monitored at holding potentials from −100 mV to +100 mV.

**Figure 2:**
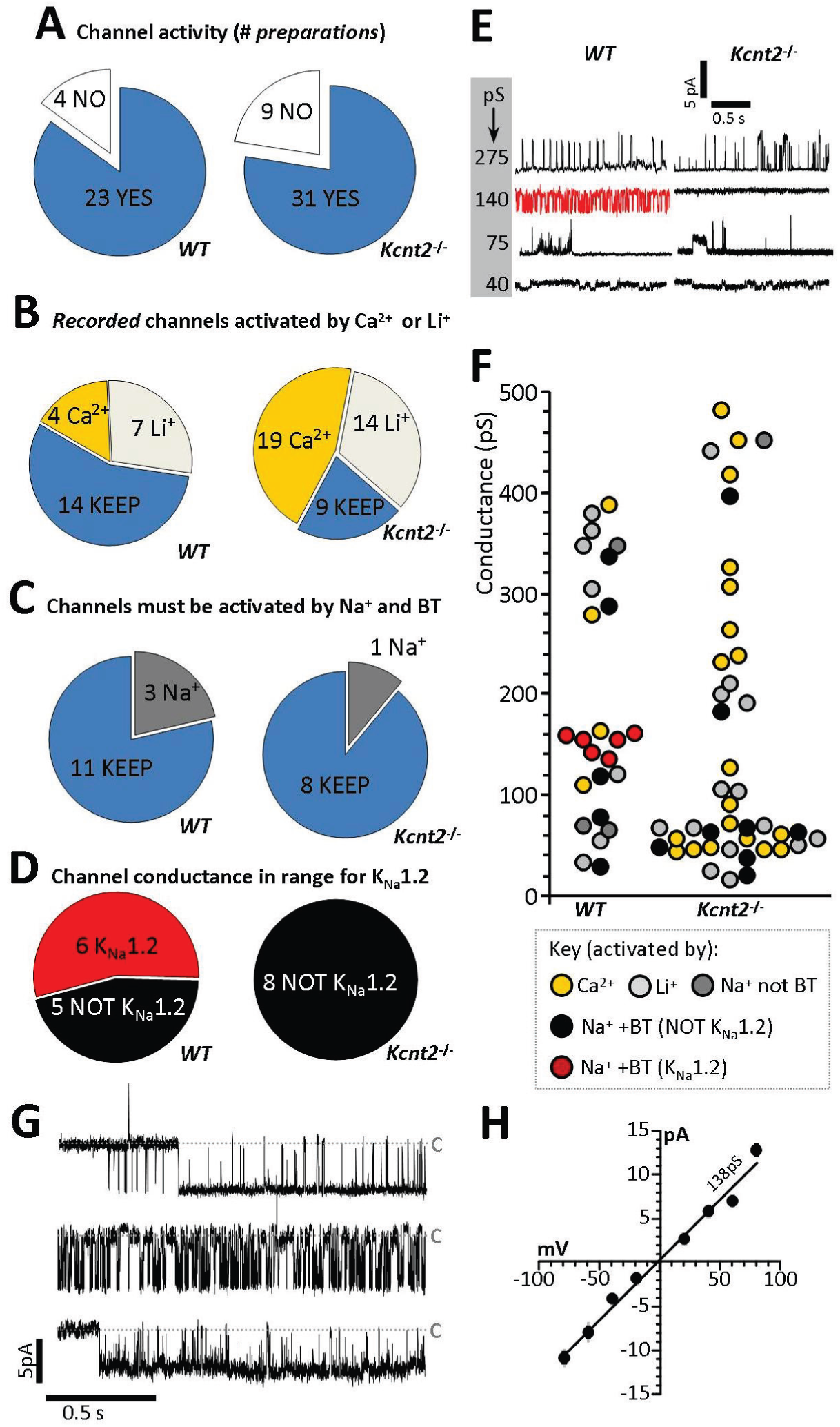
Mitochondria Contain a K_Na_1.2 Channel. **(A)** Preparations from WT (left) and *Kcnt2*^-/-^(right) mitoplasts, sorted by channels observed (blue) or no channels observed (white). **(B)** Exclusion of Ca^2+^-activated (gold) or Li^+^-activated (light gray) channels. **(C)** Selection of channels activated by Na^+^ and bithionol (BT), with exclusion of channels activated by Na^+^ alone and subsequently blocked by BT (dark gray). For panels B & C, channels carried over to the subsequent screening step are shown in blue. **(D)** Selection of channels with peak conductance matching that reported for K_Na_1.2 (red). **(E)** Example traces of channels observed in WT and *Kcnt2^-/-^* preparations with a variety of peak conductances (pS, shown in gray inset). Red trace indicates a channel assigned as K_Na_1.2. **(F)** Peak conductance of channels observed from all traces. Color key for panels B-F shown at base of panel. <u>N.B.</u> Channels with conductance >500 pS are omitted for clarity. **(G)** Example of 2 s. recordings from three K_Na_1.2 channels observed in WT mitoplasts (i.e., red points in Figure 1F) at −40 mV holding potential. Current scale bar indicated at left. Closed states are indicated by gray dashed line labeled “C”. **(H)** Channel current vs. voltage plot of peak unitary conductances of all K_Na_1.2 channels from WT mitoplast recordings. Average slope conductance was 138±1 pS.

First, patches exhibiting channel activity in Ca^2+^ alone were discarded, as this is indicative of K_Ca_ channel activity (Figure 2B, gold). Next, we discarded patches exhibiting activity following the switch to LiCl, as this is indicative of Na^+^-conductance (31) (Figure 2B, light gray). Next, patches exhibiting channel activity in the presence of activating levels of NaCl (40mM) and NaCl plus BT (2.5 µM) were considered potential K_Na_1.2 candidates (Figure 2C blue). Recordings exhibiting channel activity in the presence of Na^+^ but not following BT addition, were also discarded (Figure 2C dark gray). Finally, single channel unitary conductance was compared to expected values for K_Na_1.2 (~140 pS) (15, 31, 32) and those patches exhibiting appropriate conductance were considered to be K_Na_1.2 channels worthy of further analysis (Figure 2D red).

Figure 2E shows representative recordings observed in mitoplast patches from WT and *Kcnt2*^-/-^ mice, exhibiting a variety of unitary conductances, with the knockouts having an absence of activity in the conductance range expected for K_Na_1.2. Quantitation of all conductances showed a cluster of 6 channels in WT mitoplasts that passed all screens (Figure 2F red data points), with no channels of similar conductance observed in *Kcnt2*^-/-^ mitoplasts. The *Kcnt2*^-/-^ recordings contained proportionally more Na^+^ and BT activated channels than WT in the small conductance range (Figure 2F, black symbols in 20-80 pS range, *Kcnt2*^-/-^ 6 of 42 vs. WT 2 of 25). A similar observation has been made for a mitochondrial BK channel, wherein loss of the expected conductance in a knockout was accompanied by appearance of smaller conductances (12). Figure 2G shows representative examples of single channel activity for three (of six) Na^+^ and BT activated conductances recorded at a holding potential of −40 mV in WT mitoplasts. As previously reported for K_Na_1.2 (15), these channels showed rapid flickering between open and closed states. The average unitary slope conductance was 138±1 pS (Figure 2H: peak conductance graph for all six channels). Under our buffer conditions, reversal potential for channels conducting Na^+^ or Cl^-^ is predicted to be −23 mV or −8 mV respectively. Since K^+^ is the only other ion present, and the channel current crossed zero at −2 mV, this indicates that K^+^ is the predominant conducting ion. Together, the data in Figure 2 demonstrate that WT cardiac mitochondria contain a K^+^ channel with the ion-selectivity, ion-sensitivity, pharmacology and conductance properties of K_Na_1.2, that is absent in mitochondria from *Kcnt2*^-/-^ mice.

### Electrophysiologic Characterization of Mitochondrial K_Na_1.2

A detailed single channel analyses of all 6 assigned K_Na_1.2 patches was not possible due to the presence of two or more identical channels within some patches (Figure 3A), suggesting that K_Na_1.2 channels may cluster in their endogenous membranes. Such clustering has been previously reported for K_Na_1.1 channels in neuronal plasma-membranes (33). Examination of all suitable WT recordings (representative example Figure 3B) revealed a higher open probability at negative potentials (Figure 3C). In addition, as previously reported for K_Na_1.2 (15, 35), numerous distinct subconductances were apparent between 35 and 140pS (Figure 3D). This behavior is characteristic of K_Na_1.1 channels (*Kcnt1*/Slack/SLO2.2) (34, 35) and while it has been observed in K_Na_1.2 mutants (36), this is the first observation of such in WT K_Na_1.2 channels in endogenous membranes. The average slope conductance when considering all subconductances was 75 pS (Figure 3E), which agreed well with the average chord conductance of 74.8±6.8 pS (assuming reversal potential = 0 mV). These data indicate that smaller subconductance states predominate in the active channel current.

**Figure 3:**
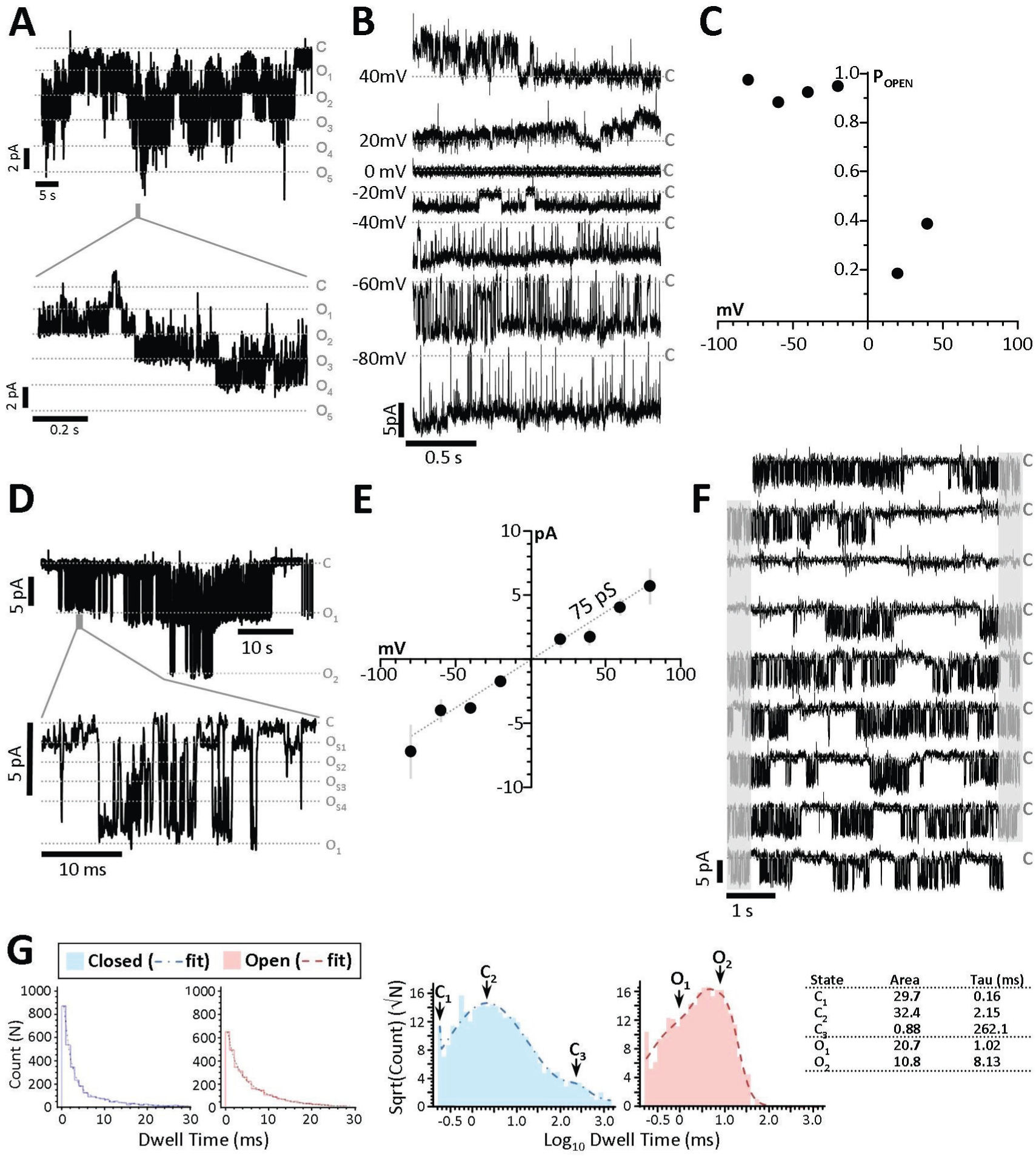
Single Channel Characteristics of Mitochondrial K_Na_1.2. **(A)** Expanded traces from recordings of patches containing five mitochondrial K_Na_1.2 channels (holding potential −20mV) and a time-expanded trace for the region highlighted by the gray bar in the trace above. Gray dotted lines represent closed (“C”) and multiple open (O_1_, O_2_, O_3_, etc.) states. **(B)** Traces from a single K_Na_1.2 channel, at holding potentials of 40mV to −80mV. **(C)** Channel open probability plot from the channel shown in panel B. **(D)** Representative trace of a recording with two channels on a compressed time scale (upper) and a time-expanded trace for the region highlighted by the gray bar in the trace above, showing multiple subconductance states within the channel peak conductance (O_S1_, O_S2_, O_S3_ etc.). **(E)**Current Voltage relationship of all six mito-K_Na_1.2 channels showing average current at each holding potential. The decreased slope conductance (compared to Figure 2H showing peak unitary conductance) indicates that subconductances averaging 75 pS dominate the average current during the recordings. **(F)** 45 s continuous trace of a single channel. Gray areas indicate portions of each trace (right) which are repeated (left) on the next line. Closed states are indicated by gray dashed line labeled “C”. **(G)** Log binned channel closed (sky blue) and open (salmon) dwell-time peaks and frequency of closed dwell times plotted against their duration. Table insert shows calculated Area and time constant() values.

Open and closed channel dwell times were also calculated from continuous recordings at −40mV (representative 45 s. example, Figure 3F). The frequency of channel open and closed dwell times was fitted to the sum of multiple simple exponentials, constituting 94.6% of the total area under the curve, and subsequent log-binned histograms revealed peaks representing two open times and three closed times (Figure 3G). The longest closed time ( _3_ = 262 ms.) represents long periods of channel closure between bursts of activity, which is characteristic for K_Na_1.2 (REF).

### K_Na_1.2 Channel Activation Uncouples Cardiomyocyte Oxidative Phosphorylation

We next sought to determine the impact of cardiac K_Na_1.2 channel activity on cardiomyocyte bioenergetics. Using Seahorse^™^ extracellular flux (XF) analysis, we measured oxygen consumption rates (OCR) of cardiomyocytes from WT and *Kcnt2*^-/-^ mice. Cells from both genotypes had similar viability and rod-shaped morphology (Figure 4A). Figure 4B shows that the K_Na_ opener BT (2.5 µM) significantly stimulated OCR in oligomycin-treated cardiomyocytes from WT mice but not those from *Kcnt2*^-/-^ mice (WT: 370±48 Max OCR; *Kcnt2*^-/-^ 154±32 Max OCR). The small effect of BT seen in *Kcnt2*^-/-^ cells is consistent with the observation of the small conductances activated by Na^+^ and BT in *Kcnt2*^-/-^ mitoplast patches (Figure 2F).

**Figure 4:**
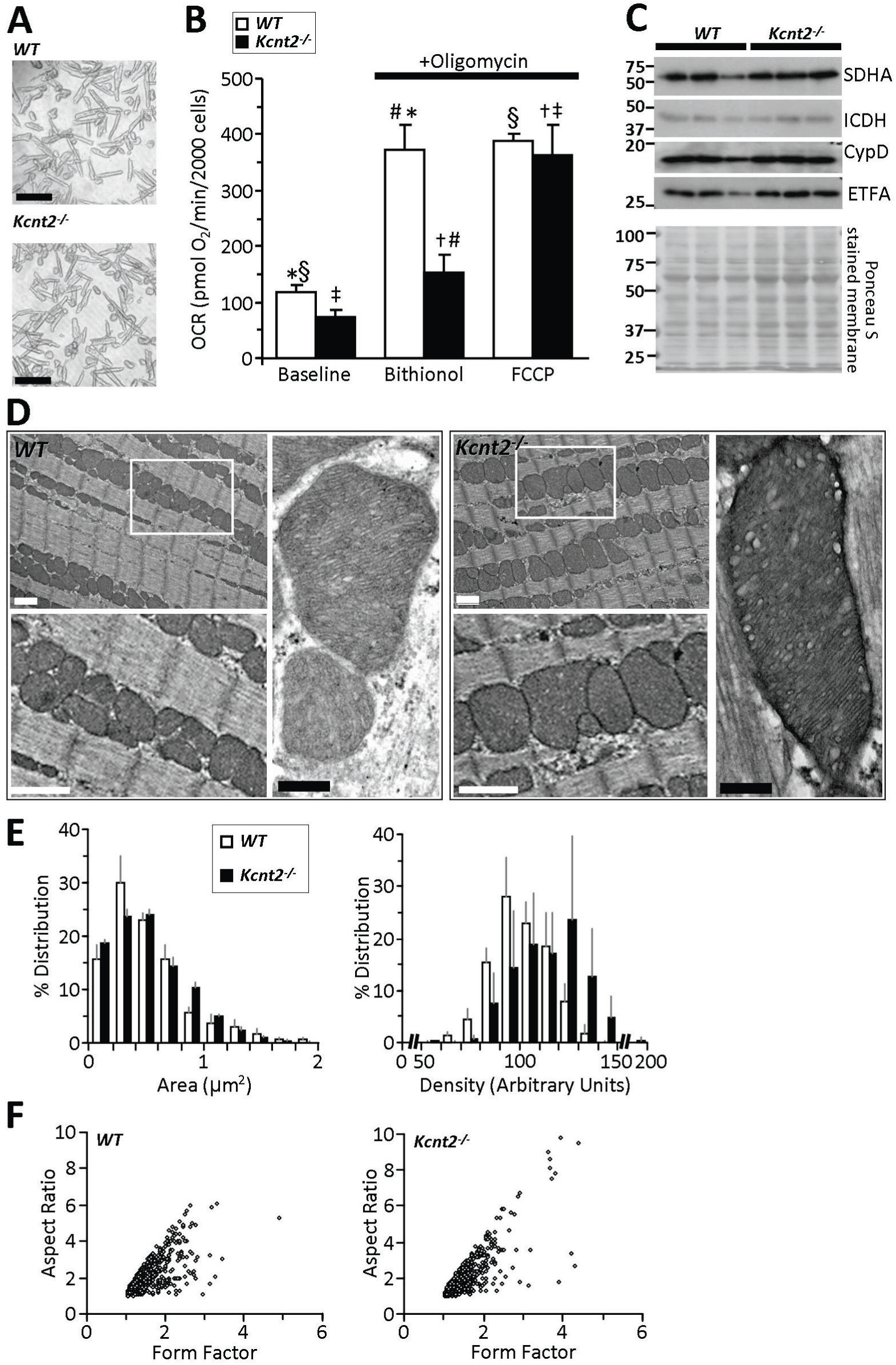
Cardiomyocyte Bioenergetics and Mitochondrial Ultrastructure. **(A)** Representative images of isolated cardiomyoctes from WT and *Kcnt2^-/-^* hearts. Black scale bar is 100 µm. **(B)** Oxygen consumption rate (OCR) of isolated cardiomyocytes measured in XF96 Seahorse Analyzer, with addition of oligomycin (1µg/ml) and either 2.5 µM bithionol (K_Na_ opener) or 500 nM FCCP (mitochondrial uncoupler). Statistics were measured using 2-way ANOVA with Bonferroni correction and post-hoc *t*-test. Bars with the same symbol are significantly different from each other (p<0.05). Data are means±SEM, N=4-5. **(C)** Western blots from WT and *Kcnt2^-/-^* heart homogenates showing levels of mitochondrial proteins (SDHA, ICDH, CypD and EFTA), and Ponceau stain loading control. **(D)** Representative transmission electron microscope (EM) images from fixed heart slices. For each genotype, lower left panels show inset boxes at higher magnification. Both white scale bars = 1 µm. Right panels show increased magnification of single mitochondria from WT and *Kcnt2*^-/-^ mice with mitochondrial ultrastructure visible (i.e., cristae folds, outer and inner membrane contacts). Black scale bar 200 nm. **(E)** Binned histogram of mitochondrial area or mitochondrial density, obtained from analysis of EM images using ImageJ software. Data are means±SEM for each bin, N=3-4. **(F)** Form-factor/aspect-ratio scatter plot. Values in panels E and F were obtained from N=1054/789 mitochondria, from 17/14 fields of view, from 4/3 hearts, of WT/*Kcnt2^-/-^*.

In the mitochondrial K^+^ cycle (37), K^+^ entry to the organelle activates a mitochondrial K^+^/H^+^ exchanger, such that mitochondrial K^+^ channel activity can uncouple oxidative phosphorylation. As such, the much larger BT induced respiratory stimulation in WT vs. *Kcnt2*^-/-^ cardiomyocytes is likely due to mitochondrial uncoupling. As an additional control, the bona-fide mitochondrial uncoupler FCCP elicited similar maximal respiration rates in cardiomyocytes from both genotypes WT: 387±15 Max OCR; *Kcnt2*^-/-^ 364.2±53 Max OCR), rendering it unlikely that the differential effect of BT in WT vs. *Kcnt2*^-/-^ cells was due to an underlying difference in overall bioenergetic capacity. Consistent with this, western blotting for a number of mitochondrial marker enzymes (SDHA, ICDH, Cyp-D, and ETFA) suggested no difference in mitochondrial mass or content between WT and *Kcnt2*^-/-^ hearts (Figure 4C).

### Loss of K_Na_1.2 Mildly Impacts Cardiac Mitochondrial Ultrastructure

An important function of the mitochondrial K^+^ cycle is the regulation of organelle volume (38, 39), and K_Na_1.2 activity is known to be sensitive to osmolarity (40, 41). Thus we hypothesized that mitochondria from *Kcnt2*^-/-^ may exhibit ultrastructural changes. Electron-microscopic analysis of hearts from WT and *Kcnt2*^-/-^ mice (Figure 4D-F) revealed that mitochondria had similar area (WT: 0.51±0.30 µm^2^, *Kcnt2*^-/-^: 0.53±0.32 µm^2^, means±SD, N=3-4) and matrix density (WT: 109±13, *Kcnt2*^-/-^: 102±11, means±SD, N=3-4). However, the distribution of these parameters (Figure 4E) suggests a small shift toward increased area and density in *Kcnt2*^-/-^vs. WT. Additionally, no difference in either form-factor or aspect-ratio was observed between genotypes (Figure 4F), indicating that K_Na_1.2 deficiency does not alter mitochondrial fission or fusion (42, 43). Together, the data in Figure 4 suggest that although activation of K_Na_1.2 can uncouple respiration, loss of the channel does not impact cardiac mitochondrial structure or content.

### K_Na_1.2 is Required for Cardiac Respiratory Reserve Capacity when Oxidizing Fat

In an effort to further understand the bioenergetic effects of K_Na_1.2 deficiency we compared metabolic substrate preferences in cardiomyocytes isolated from WT and *Kcnt2*^-/-^ mice. Myocytes were incubated with either: (i) glucose alone (with etomoxir to inhibit fatty acid - oxidation), (ii) palmitate alone (with 2-deoxyglucose to inhibit glycolysis), or (iii) glucose plus palmitate. The response to uncoupling by FCCP (500 nM) was used to determine “respiratory reserve” (RR) capacity under each substrate condition (Figures 5A-C).

**Figure 5:**
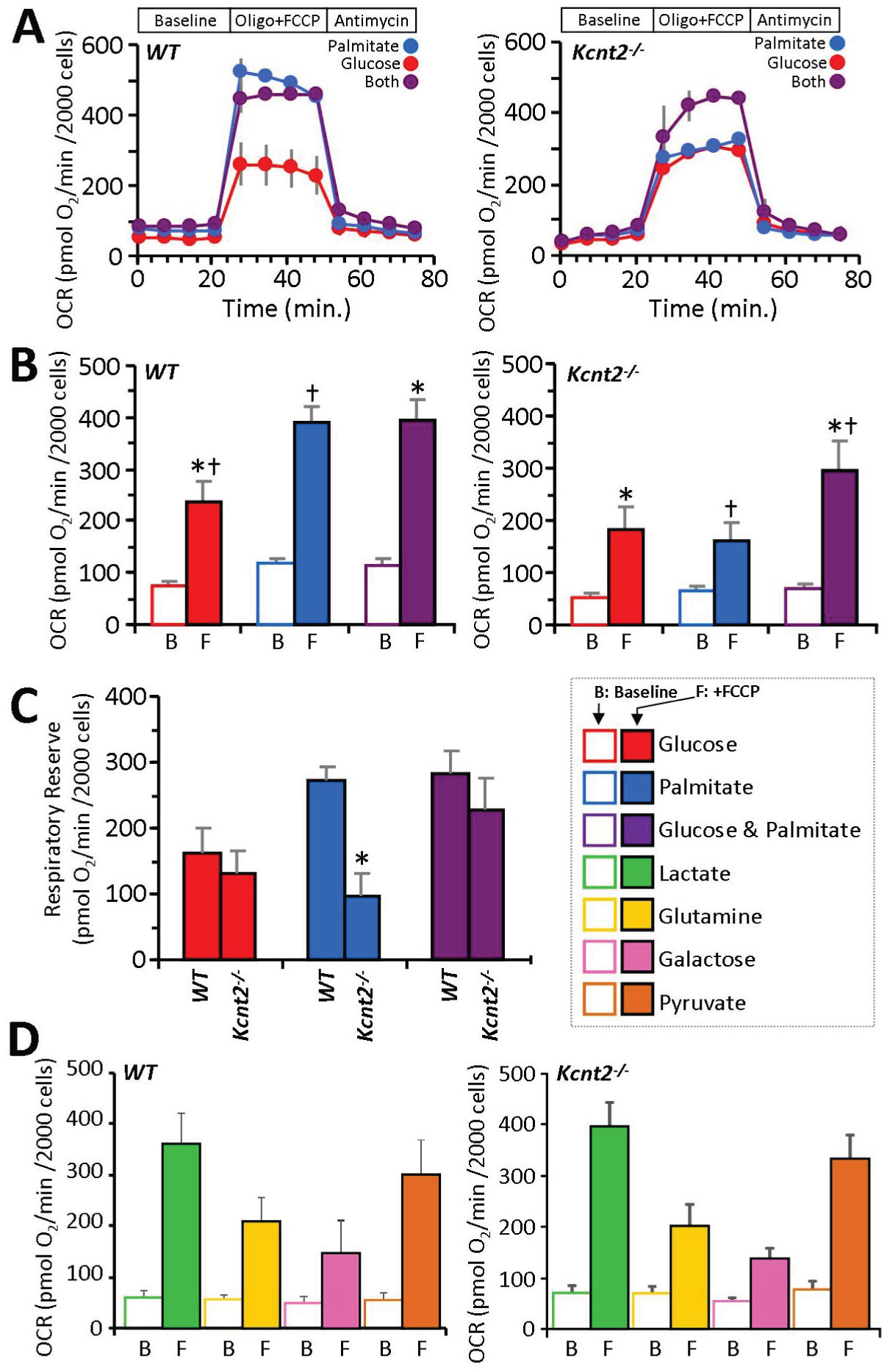
K_Na_1.2 Loss Impacts Cardiac Metabolic Substrate Choice under Stress. **(A)** Representative Seahorse XF oxygen consumption rate (OCR) traces of isolated WT or *Kcnt2^-/-^* cardiomyocytes, incubated in the presence of different metabolic substrates: glucose (red), palmitate (blue), glucose + palmitate (purple). Data from a single XF plate are shown (means ± SD of 12 wells per substrate or genotype). Timeline above traces shows OCR was measured at baseline, then with addition of the ATP synthase inhibitor oligomycin (1 µg/ml) plus the mitochondrial uncoupler FCCP (500 nM), and finally with the mitochondrial complex III inhibitor antimycin A (5 µM). **(B)** Group OCR averages of baseline (B) and FCCP-uncoupled (F) cardiomyocytes under substrate conditions as defined above. Statistics were measured using 2-way ANOVA with Bonferroni correction and post-hoc *t*-test. Data are means ± SEM, for N=4 *Kcnt2^-/-^* or 5 WT, independent cardiomyocyte preparations. Bars with the same symbol are significantly different from each other (p<0.05). **(C)** Respiratory reserve (RR) capacity calculated from the data in panel B (i.e., uncoupled minus baseline OCR). Means ± SEM, N=4-5, ^*^p<0.05 between genotypes. Color key for metabolic substrates used in all panels is shown to the right of panel C. **(D)** Oxygen consumption rates (OCR) of WT and *Kcnt2^-/-^* cardiomyocytes metabolizing different substrates: lactate (lime), glutamine (gold), galactose (rose), or pyruvate (copper). Empty bars = baseline (B), Filled bars = FCCP uncoupled (F). Data are means ± SEM from 4-5 independent cardiomyocyte preparations.

Cardiomyocytes from WT and *Kcnt2*^-/-^ cells exhibited a similar baseline OCR under all substrate conditions (Figure 5B open bars). In WT cells, a robust uncoupling response to FCCP was seen under all conditions, and notably the uncoupling response with palmitate alone (3.3 fold) was equal to that seen when both substrates were present (3.3 fold). However, in *Kcnt2*^-/-^ cells, the uncoupling response with palmitate alone (2.4 fold) was significantly blunted compared to that seen when both substrates were present (4.3 fold) (comparison between blue and purple bars in left & right panels of Figure 5B). The additional OCR induced over baseline by FCCP is used to calculate RR capacity. Figure 5C shows that the RR of WT and *Kcnt2*^-/-^ cells is similar in either the glucose alone or the glucose plus palmitate conditions (red and purple bars respectively). However, with palmitate alone (blue bars), *Kcnt2*^-/-^ cells exhibit a significant RR deficit relative to WT. Notably, no such RR deficit was observed in myocytes from *Kcnt2*^-/-^ mice respiring on lactate, glutamine, galactose, or pyruvate (Figure 5D), indicating the *Kcnt2*^-/-^ RR deficit is specific to fat oxidation.

To test the physiologic relevance of this RR deficit, the ability of perfused hearts to respond to increased metabolic demand was tested. Hearts from WT and *Kcnt2*^-/-^ mice were perfused with palmitate as the sole carbon source, while stimulating workload by addition of the - adrenergic agonist isoproterenol (100nM). Hearts from *Kcnt2*^-/-^ mice showed a significantly reduced functional response to isoproterenol, relative to WT (WT: 213±20 % vs. *Kcnt2*^-/-^: 159±13 %, means±SEM, N=7) (Figure 6A). However, consistent with the isolated cardiomyocyte OCR data (Figure 5C), no difference in the isoproterenol-induced functional response was observed when the perfusion buffer was supplemented with glucose and palmitate (Figure 6A). Together, these data suggest that loss of K_Na_1.2 results in an impaired ability to respond to increased metabolic demand when oxidizing only fat. Consistent with previous reports (3, 18) no EKG differences were observed in *Kcnt2*^-/-^mice (Figure 6B), suggesting that loss of K_Na_1.2 *per se* does not impact cardiac contractile function.

**Figure 6:**
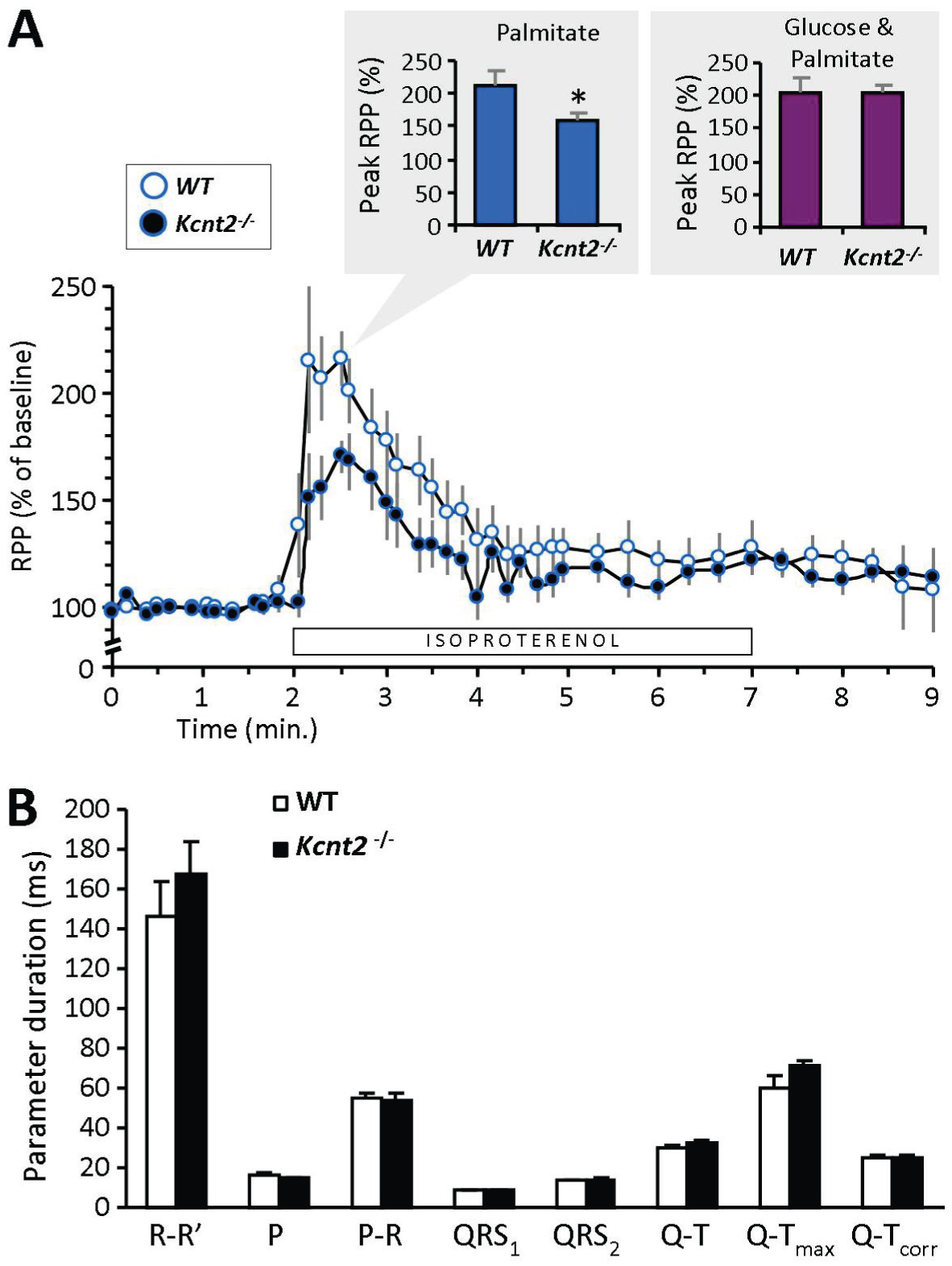
**(A)** Cardiac function data (heart rate × pressure product, RPP) for isolated perfused hearts from WT or *Kcnt2*^-/-^ mice, perfused with Krebs-Henseleit (KH) buffer containing palmitate as the sole metabolic substrate. Bar below the trace indicates duration of 100 nM isoproterenol infusion. Graph shows RPP as a % of the average baseline value for 1 min before isoproterenol infusion. Inset bar graph shows comparison of the peak response to isoproterenol under this substrate condition (palmitate only, blue). Adjacent inset (right) shows the peak response to isoproterenol from a separate series of perfusions in which the KH buffer contained both palmitate and glucose as substrates (purple). Data are means ± SEM, N=7, ^*^p<0.05 between genotypes. **(B)** EKG parameters obtained *in-vivo* from avertin-anesthetized WT and *Kcnt2^-/-^* mice. R-R’ = distance between R waves of each beat (i.e., 1/HR), P = p-wave duration. P-R = interval between P and R waves. QRS_1_ & QRS_2_ = diameter of QRS complex (different calculation algorithms). Q-T = interval between Q and peak of T wave. Q-T_max_ = interval between Q and end of T wave. Q-T_corr_ = QT interval corrected for heart rate). Data are ± SD, N=4-5 animals.

### Whole Animal Metabolic Differences in Kcnt2^-/-^ Mice

Since the heart is an major fat-burning organ, we hypothesized that the fat-specific RR capacity deficit in *Kcnt2*^-/-^ might be accompanied by metabolic perturbations at the whole animal level. No significant differences in weight gain were observed between WT and *Kcnt2*^-/-^ mice over 25 weeks (Figure 7A). Analysis of percent body fat content by differential energy X-ray absorptometry analysis (DEXA, Figure 7B) revealed a small difference in average fat content between genotypes (WT: 12.1±2.8 % vs. *Kcnt2*^-/-^: 13.6±3.3 %), but nevertheless this difference was statistically significant between paired littermates (Figure 7C). In addition, while WT mice showed an expected drop in blood glucose following an overnight (15 hr.) fast, no such drop was seen in *Kcnt2*^-/-^ mice (Figure 7D). This suggests elevated gluconeogenesis in response to fasting in *Kcnt2*^-/-^, which is consistent with a potential increased reliance on glucose oxidation as a compensatory response to the fat-specific RR defect under stress conditions.

**Figure 7:**
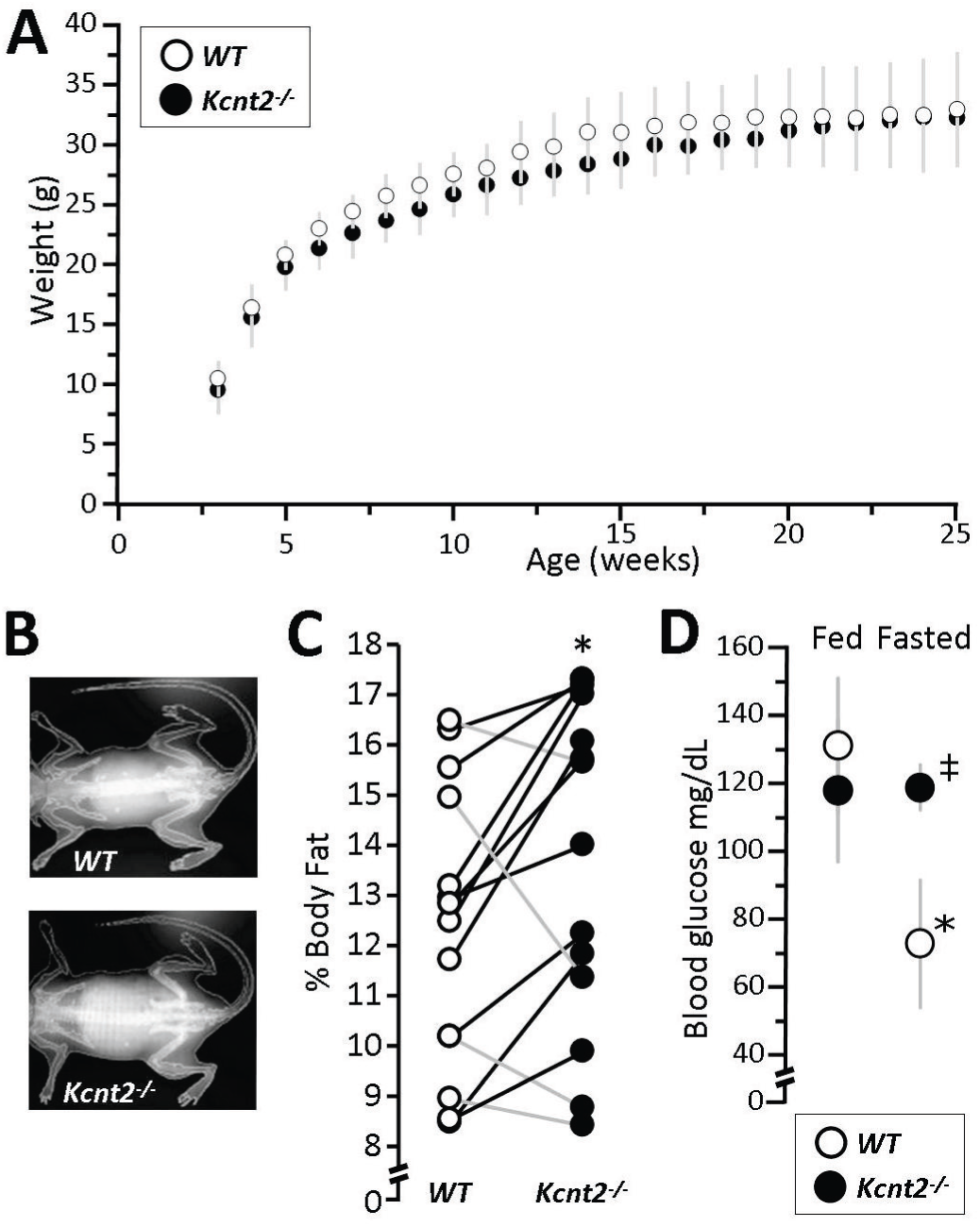
Loss of K_Na_1.2 Impacts Whole Body Metabolic Phenotype. **(A)** Body weights of WT and *Kcnt2^-/-^*mice from weaning (3 weeks) to 25 weeks of age. Data are means ± SD, N=12. **(B)** Representative DEXA images from WT and *Kcnt2^-/-^* mice. **(C)** Percent body fat measured by DEXA scan of WT (white symbols) and *Kcnt2^-/-^* (black symbols) littermates (pairs are indicated as data points connected by lines), N=14. ^*^p<0.05 between genotypes by paired *t*-test. **(D)** Blood glucose levels measured in WT and *Kcnt2*^-/-^ mice at baseline (5 PM, Fed) and following a 15 hr fast (8 AM, Fasted). Data are means ± SD, N=3. ^*^p<0.05 between fed and fasted state within a genotype. ‡p<0.05 between genotypes at the same time point.

### Metabolomic and Expression Profiling of Kcnt2^-/-^ Hearts

To investigate the molecular underpinnings of the fat-specific RR defect in *Kcnt2*^-/-^ hearts, expression of metabolic regulatory genes was measured using a predesigned qPCR array (Figure 8A, Table 1). No differences were observed between WT and *Kcnt2*^-/-^ hearts, suggesting the fat-specific RR defect is not due to a remodeling of metabolism at the gene level. Separately, a small but non-significant decrease in energy charge (ATP+½ADP/(ATP+ADP+AMP)) was observed in *Kcnt2*^-/-^ mice (WT: 0.87±0.05 vs. *Kcnt2*^-/-^: 0.77±0.03, mean±SEM, N=5-6), suggesting that energy-sensing metabolic regulators such as AMP dependent protein kinase (AMPK) may be altered. However, western blotting analysis revealed no difference in AMPK phosphorylation between WT and *Kcnt2*^-/-^ hearts (data not shown). Together with data in Figures 4 and 5, these findings suggest that K_Na_1.2 deficiency does not induce large scale remodeling of cardiac mitochondrial metabolism. Rather, K_Na_1.2 loss specifically impacts cardiac fat oxidation, only under conditions of high energy demand such as uncoupling or - adrenergic stimulation.

**Figure 8:**
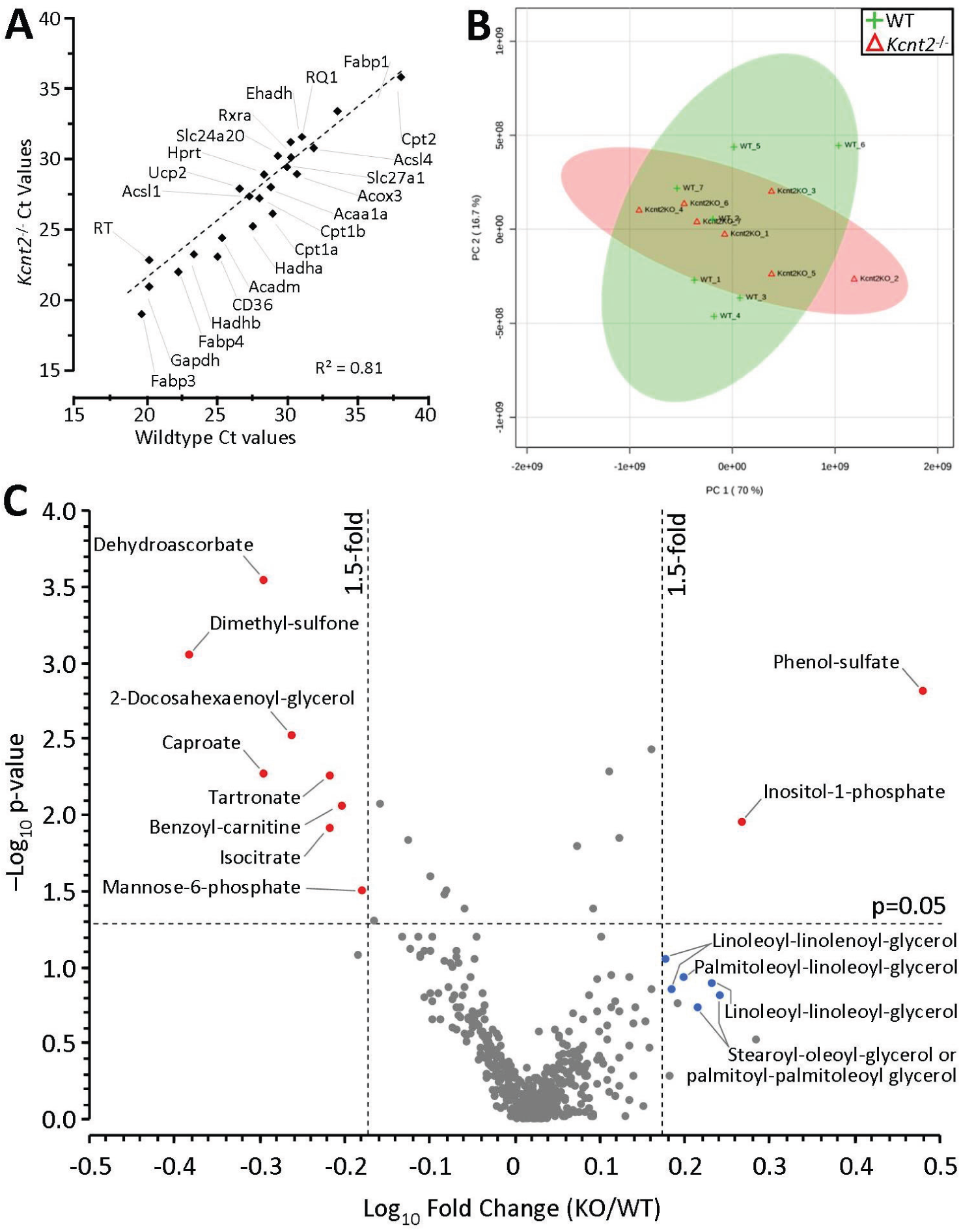
Cardiac Expression Profiling and Metabolomics. **(A)** qPCR Ct values for 27 metabolically important genes (see Table 1), in WT and *Kcnt2*^-/-^ hearts. N=3 independent RNA preparations per genotype. Data are means, errors are omitted for clarity. **(B)** WT and *Kcnt2*^-/-^ hearts were perfused with KH buffer containing palmitate plus glucose, and freeze-clamped for metabolomic analysis by LC-MS/MS. Graph shows principle component analysis of 501 cardiac metabolites. The first and second principal components contributed 86.7% of the overall metabolic character. Shaded ovals overlaying the graph indicate 95% confidence intervals for WT (lime) and *Kcnt2*^-/-^ (salmon pink) samples. **(C)** Volcano plot of the metabolic profile of *Kcnt2*^-/-^ vs. WT hearts. Axes show - Log_10_(p-value) vs. Log_10_(fold change). Dashed lines show p=0.05 cut off (y axis) and 1.5fold change cut off (x-axis). Each point represents a single metabolite, and data for each point are means from N=7 hearts. Errors are omitted for clarity. Metabolites passing fold-change and p-value criteria are highlighted red. Additional metabolites discussed in the text are highlighted blue.

**Table 1.**
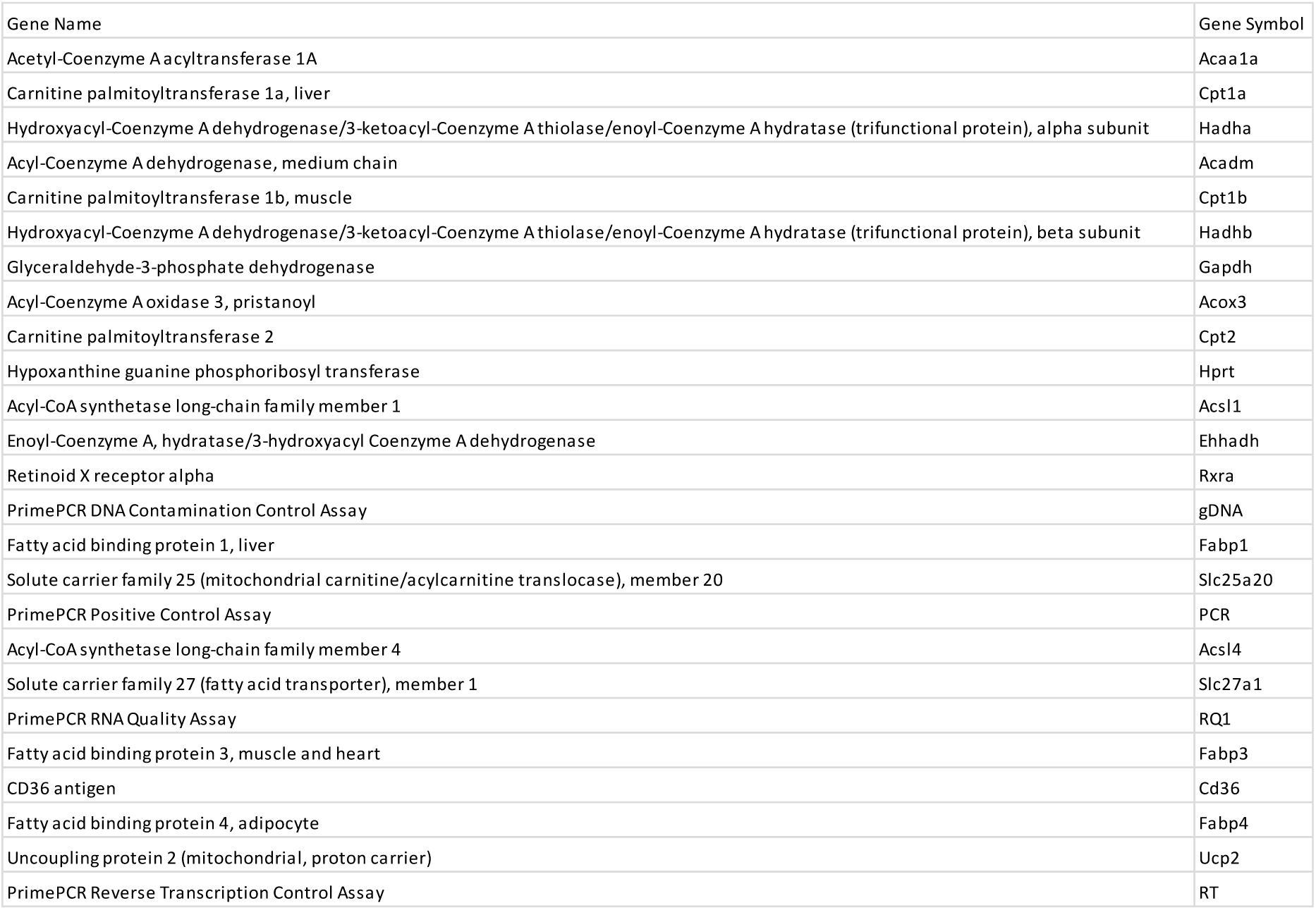
Gene names and symbols for genes targeted by BioRad PrimePCR qPCR assay kit (Figure 8A)

Finally, to understand the effects of K_Na_1.2 deficiency on cardiac metabolism at a systems level, an unbiased metabolomics analysis was performed. Principal component analysis (PCA) showed no significant difference in the fundamental character of metabolism between WT and *Kcnt2*^-/-^hearts at baseline (Figure 8B). A volcano plot for all 501 metabolites measured revealed only 10 were significantly altered (>1.5-fold vs. WT, p<0.05) (Figure 8C). Of these metabolites, notable changes were an increase in phenol-sulfate and decrease in dimethyl-sulfone, potentially indicating perturbations in aryl-sulfotransferase activity and sulfur metabolism. Dehydroascorbate was significantly lower in *Kcnt2*^-/-^ hearts, potentially indicating lower oxidative load. In addition, inositol-1-phosphate was significantly elevated, and a cluster of diacylglycerol metabolites was also elevated (although no individual DAG approached significance), suggesting enhanced phospholipase C activity in *Kcnt2*^-/-^. Overall, the comparatively minor nature of metabolomic perturbations in *Kcnt2*^-/-^ hearts at baseline is consistent with the notion that the impact of K_Na_1.2 loss is limited to fat oxidation under conditions of high energy demand.

## Discussion

A plethora of studies has identified mitochondrial K^+^ channels at the phenomenological level (reviewed in (1, 44)), and several studies have linked these channels mechanistically to protection against IR injury (1, 45- 47). However, surprisingly few examples exist of bona-fide mitochondrial K^+^ channels that are: (i) identified at the molecular (genetic) level, (ii) characterized with robust electrophysiology, and (iii) linked to any specific mitochondrial channel function phenotype (examples include K_Ca_1.1 (11, 12), K_ATP_ (10), and SK3 (48)). While the phenomenon of a K_Na_ channel was first characterized in hearts thirty years ago, the field of K_Na_ research rapidly transitioned to study of these channels in brain (19). As such, very little is known about role of K_Na_ in cardiomyocytes, including sub-cellular localization. Using patch-clamp studies of isolated cardiac mitochondrial inner membranes, we recorded a K^+^ channel matching the known characteristics of K_Na_1.2 (ion-sensitivity, ion-selectivity, pharmacology and conductance) in mitoplasts from WT mice, that was absent in those from *Kcnt2*^-/-^ mice (Figures 1-3). This is the first report of a K_Na_ channel in mitochondria.

Since cardiac mitochondria are reported to contain several cation and anion channels (49, 50), mitoplast recordings were subjected to a rigorous screening process to ensure that conductances assigned as K_Na_1.2 did not originate from other channels. Bath/pipette ratios of the major ion components (K^+^, Na^+^, and Cl^-^) enabled reversal potentials to be differentiated. The Na^+^ surrogate Li^+^ was used to screen out any Na^+^-activated currents that were in fact due to Na^+^ conductance. In addition, any conductances activated by Ca^2+^ were dismissed, to ensure patched membranes were free of K_Ca_ channels. Anion channels of the IMAC and CLIC variety exhibit conductances (~100pS: IMAC, ~8pS: CLIC4) sufficiently distinct from K_Na_1.2 that, in combination with the observed reversal potential, we are confident do not contribute to assigned K_Na_ conductances. Similarly mitoK_ATP_ channels (~10pS) would not be mistaken for K_Na_1.2. Finally, K_Na_1.2 activation is significantly enhanced by Cl^-^ in the presence of activating Na^+^, hence Cl^-^ salts were used to maximize the probability of detecting these channels (15). Of 25 recordings in WT mitoplasts, this strategy yielded 6 channels, a 24% success rate. Application of this rate to the 42 recordings in *Kcnt2^-/-^* mitoplasts predicts 10 such channels, but we observed zero.

Several known electrophysiologic characteristics of K_Na_1.2 channels were also observed in our assigned K_Na_1.2 mitoplast recordings, including: (i) multiple sub-conductance states (Figure 3D) (15, 36), (ii) rapid flickering between open and closed states (Figure 2G) (15), (iii) burst operation with prolonged closed times between bursts (Figure 3F/G) (15, 51), and (iv) multiple channels within the same patch suggesting channel clustering in membranes (Figure 3A) (36). In combination with the absence of such conductances in *Kcnt2*^-/-^mitoplasts, these characteristics strengthen the conclusion that the genetic origin of the channels herein assigned as mito-K_Na_1.2, is the *Kcnt2* gene product. Whole mitoplast attached patch experiments (Figure 1E) suggest that the orientation of the channel in the mitochondrial inner membrane is the same as that reported for plasma membrane K_Na_1.2, namely with the c-terminal Na^+^ sensing site on the inside of the membrane (15, 52).

Patches from WT and *Kcnt2*^-/-^ recordings contained channels with a sim**i**lar overall range of conductances. However, the distribution of these conductances between genotypes was shifted. In particular, channels with unitary conductances in the range of 20-80pS were more frequently observed in *Kcnt2*^-/-^ (Figure 2F). A similar observation was recently made using a cardiac-specific knockout of the mito-BK channel (12), raising the intriguing possibility that loss of one mitochondrial K^+^ channel may lead to compensatory up-regulation of other channels, to maintain K^+^ homeostasis. Given the importance of the mitochondrial K^+^ cycle for the regulation of organelle volume (37), such compensatory K^+^ fluxes may account for the persistence of a small but non-significant respiratory uncoupling effect of BT in *Kcnt2*^-/-^ cardiomyocytes (Figure 4B), as well as the relatively minor effect of K_Na_1.2 loss on mitochondrial ultrastructure (Figure 4D-F). An alternative to compensatory expression between different mitochondrial K^+^ channels, is the possibility they may form heteromers. Indeed, individual K_Na_1.1 (*Kcnt1*) subunits can combine with K_Ca_1.1 (*Kcnma1*) subunits to form functional heterotetramers of intermediate conductance and activation properties (14). Heterotetramers of K_Na_1.1 with K_Na_1.2 have also been demonstrated (36). However, the existence of K_Ca_1.1/K_Na_1.2 heterotetramers has not been reported.

A key aspect of these studies was the discovery that loss of cardiac K_Na_1.2 results in a unique metabolic phenotype, namely a specific defect in cardiac fat oxidation only under conditions of high energy demand. An important caveat is that we cannot currently rule out the possibility that the metabolic phenotype of *Kcnt2*^-/-^ is due to loss of the channel at the plasma membrane. However, evidence for the existence of K_Na_ channels at the cardiac plasma membrane comprises a single study 30 years ago (19). Despite the development of anti-K_Na_ antibodies, fluorescent tags, and K_Na_ channel knockout animals, confirmation of the channel’s existence at the cardiac plasma membrane has not been confirmed using any of these methods during the intervening time period. Coupled with definitive data herein demonstrating existence of K_Na_1.2 in mitochondria, and previous reports of a BT activated mitochondrial K^+^ channel that is absent in mitochondria isolated from *Kcnt2*^-/-^, we assert that the most likely explanation for a mitochondrial/metabolic phenotype in *Kcnt2*^-/-^, is loss of the channel in mitochondria.

The metabolic phenotype of *Kcnt2*^-/-^ (Figures 5-6) was manifest under conditions corresponding to classical bioenergetic “state 3”, wherein the mitochondrial membrane potential is consumed to generate ATP. As such, if the *in-situ* reversal potential of mitochondrial K_Na_1.2 is above zero, then channel opening could readily occur under conditions of high energetic demand when potential is lowered. In addition, the mitochondrial membrane potential is known to “flicker” *in-vivo*(53), such that transient depolarization events may activate mito-K_Na_1.2. These properties may render mitochondrial K_Na_1.2 function important under conditions of stress, such as tissue ischemia. Given the previously reported requirement of K_Na_1.2 for cardioprotection by APC (3), it is notable that the mitochondrial membrane potential depolarizes and cytosolic Na^+^ levels also rise during ischemia (54, 55). Together, these observations suggest that mito-K_Na_1.2 channels may be activated by acute perturbations in mitochondrial energy demand or under stress conditions such as ischemia. Such properties may explain why the metabolic phenotype of *Kcnt2*^-/-^ is only evident under stress.

The data in Figure 7 revealed additional chronic effects of K_Na_1.2 loss on bioenergetics that resulted in altered body fat content and fasting glucose metabolism. Due to the presence of a mitochondrial K^+^/H^+^ exchanger (KHE), activation of a mitochondrial K^+^ channel would be expected to decrease the mitochondrial pH, thus uncoupling oxidative phosphorylation and stimulating OCR, as shown in Figure 4B. As such, mild mitochondrial uncoupling by mito-K_Na_1.2 channel activators may represent a novel therapeutic avenue for obesity/diabetes/metabolic-syndrome. It is therefore notable that the anti-helminthic drug niclosamide, which activates K_Na_ channels (56), has long been known to uncouple mitochondria (57), and was recently shown to confer benefits in a mouse high-fat diet model of diabetes (58). Furthermore, a recent case report highlighted a patient with a K_Na_1.2 mutation (Q_270_E) suffering from migrating focal seizures that were non-responsive to the ketogenic diet typically used to treat such symptoms (59). Cardiac metabolomics also revealed a potential up-regulation of phospholipase C (PLC) signaling in the *Kcnt2*^-/-^ heart (Figure 8C). Since K_Na_1.2 channels are known to interact with the PLC substrate PIP_2_ (60), this raises the possibility that loss of K_Na_1.2 results in perturbation of PIP_2_/PLC signaling. An important PLC downstream target is protein kinase C epsilon (PKC), which is known to play a role in development of insulin resistance in response to a high fat diet (61). As such, in addition to mitochondrial uncoupling, mito-K_Na_1.2 channel activators may confer metabolic benefits via a PIP_2_/PLC/PKC signaling axis. A deeper investigation of the relationship between mito-K_Na_1.2 activity and metabolic regulation is thus warranted.

## Acknowledgments

We thank Christopher Lingle (Washington University, St Louis MO) for providing founders for the *Kcnt2^-/-^* mice, and Kathleen Kinally (New York University, Emeritus) for technical support in performing mitochondrial patch clamp experiments. We also thank Dana Godfrey (URMC musculoskeletal center) for support with DEXA analyses, and Karen Bentley (URMC electron microscopy core).

## Author Contributions

Charles O. Smith, Paul S. Brookes, and Keith Nehrke designed research. Charles O. Smith, Yves T. Wang, Sergiy M. Nadtochiy, and James H. Miller performed experiments. Yves T. Wang developed software for analysis of Langendorff traces. Charles O. Smith analyzed data. Charles O. Smith, and Paul S. Brookes wrote paper. Elizabeth A. Jonas, and Robert T. Dirksen contributed new analytical tools, training, and critical resources. This work was funded by grants from the National Institutes of Health: R01-GM087483 (to Paul S. Brookes, and Keith Nehrke), R01-HL071158 (to Paul S. Brookes) and R01-AR-059646 (to Robert, T. Dirksen).

## Non-standard abbreviations

IR injury: Ischemia-Reperfusion injury
IPC: Ischemic Preconditioning
APC: Anesthetic Preconditioning
K_Na_: Sodium activated potassium channel
K_Na_1.1: (channel encoded by *Kcnt1* (formerly Slo2.2)), aka Slack, K_Ca_4.1, SLO2.2
K_Na_1.2: (channel encoded by *Kcnt2* (formerly Slo2.1)), aka Slick, K_Ca_4.2, SLO2.1
BT: Bithionol, aka Bis(2-hydroxy-3,5-dichlorophenyl)Sulfide
OCR: Oxygen consumption rate
ROS: Reactive oxygen species

